# Forecasting dynamics of a recolonizing wolf population under different management strategies

**DOI:** 10.1101/2023.03.23.534018

**Authors:** Lisanne S. Petracca, Beth Gardner, Benjamin T. Maletzke, Sarah J. Converse

## Abstract

Species recovery can be influenced by a wide variety of factors, such that predicting the spatiotemporal dynamics of recovering species can be exceedingly difficult. These predictions, however, are valuable for decision makers tasked with managing species and determining their legal status. We applied a novel spatially explicit projection model to estimate population viability of gray wolves (*Canis lupus*) from 2021-2070 in Washington State, USA, where wolves have been naturally recolonizing since the establishment of the first resident pack in 2008. Using this model, we predicted the effects of 12 scenarios of interest relating to management actions (e.g., lethal removals, translocation, harvest) and system uncertainties (e.g., immigration from out of state, disease) on the probability of meeting Washington’s wolf recovery goals, along with other metrics related to population status. Population recovery was defined under Washington’s Wolf Conservation and Management Plan as four breeding pairs in each of three recovery regions and three additional breeding pairs anywhere in the state. The baseline, translocation, and 50% immigration scenarios indicated a high (>60%) probability of wolf recovery in Washington over the next 50 years, but scenarios related to harvest mortality (removal of 5% of the population every six months), increased lethal removals (removal of 30% of the population every four years), and cessation of immigration from out of state resulted in low probabilities (0.07, 0.12, and 0.12, respectively) of meeting recovery goals across all years (2021-2070). All but one management scenarios exhibited a geometric mean of population growth that was ≥1, indicating long-term population stability or growth, depending on the scenario. Our results suggest that wolves will continue to recolonize Washington and that recovery goals will be met so long as harvest and lethal removals are not at unsustainable levels and adjacent populations support immigration into Washington.

## INTRODUCTION

The spatiotemporal dynamics of recovering species are challenging to predict. This challenge is compounded by the variety and complexity of factors that may influence recovering species. Factors with uncertain effects may influence habitat, including land use change (Lawler et al. 2002) and altered climate regimes (Abrahms et al. 2022), while others may affect demography directly, including interspecific competition (Elbroch et al. 2015), disease (Rhodes et al. 2011, Gordon et al. 2015), and changing rates of immigration or emigration (Lieury et al. 2016, Grauer et al. 2019). Management actions themselves can also be sources of uncertainty, as their effects on abundance and vital rates can be challenging to predict (Saunders et al. 2018). Despite the challenges of predicting spatiotemporal dynamics of recovering species, this information is often required to manage populations and to determine their legal status (e.g., Smith et al. 2018).

Gray wolves (*Canis lupus*), after more than a century of widespread eradication efforts (Mech 1970), have now returned to a large and growing part of their historical range in the upper Midwest, northern Rocky Mountains, and increasingly in the Pacific Northwest (Mech 1995, Maletzke et al. 2016, Jimenez et al. 2017). Much has been learned about the ecological factors influencing recovering wolf populations, such as high survival rates (Hayes and Harestad 2000, Smith et al. 2010), high reproductive rates (Mech 1995), immigration via long-distance dispersal (Mech 1995, Hayes and Harestad 2000, Vilà et al. 2003, Maletzke et al. 2016), and presence of large core habitat areas (Smith et al. 2010). However, wolf populations are also susceptible to human-caused mortality and disease (e.g., canine parvovirus and distemper), which can impact survival rates, dispersal rates, and thus also impact populations (Mech et al. 2008, Smith et al. 2010, Nelson et al. 2012). Habitat fragmentation and barriers to dispersal can also cause population declines and hinder recovery (Geffen et al. 2004, Stronen et al. 2012).

A variety of management actions have been applied to manage wolf populations throughout their range, both to aid in their recovery and to limit their population sizes. Reintroduction, the translocation of a species into a portion of its historical range from which it has been extirpated (Armstrong and Seddon 2008), was used to return wolves to the Greater Yellowstone Ecosystem and central Idaho (Bangs and Fritts 1996) and is now being planned in the state of Colorado (Ditmer et al. 2022). By contrast, other management tools have been applied to reduce wolf numbers and to manage human-wolf conflict, including recreational harvest (Creel et al. 2015) and lethal removals for livestock and ungulate predation concerns (Bangs et al. 2006, DeCesare et al. 2018). In any recovering carnivore population, there are likely to be potentially conflicting management objectives, including both supporting recolonization of former habitat and minimizing negative impacts to human livelihoods and other social values.

The gray wolf is a state-endangered species in Washington State, USA, and as of December 2022 was federally listed under the Endangered Species Act in the western two-thirds of the state. Washington’s wolf recovery plan identifies recovery criteria, which specify four breeding pairs in each of the three recovery regions – Eastern Washington, Northern Cascades, and Southern Cascades and Northwest Coast – with three additional breeding pairs anywhere in the state (Wiles et al. 2011) to begin the process for delisting. To date, recovery goals had been met in the first two recovery regions but there were no breeding pairs in the Southern Cascades and Northwest Coast. Maletzke et al. (2016) implemented a projection model and predicted that, with immigration from British Columbia and Idaho, recovery goals were likely to be met by 2021; however, their model may have overpredicted demographic and pack size parameters, as they only had access to data from wolf populations in nearby states (Idaho, Montana, and Wyoming) and made simplifying assumptions about Washington wolf dynamics based on those data. Their work highlights the importance of capturing wolf population dynamics that are specific to Washington, including mechanisms of dispersal. The status of wolves in Washington, along with the potential for increasing human-wildlife conflicts, has created a high priority need for predicting the spatiotemporal dynamics of Washington’s wolf population under a suite of possible future scenarios.

We applied a novel spatially explicit projection model described in Petracca et al. (2024) to conduct a population viability analysis (Morris and Doak 2002) for wolves in Washington State. We considered 12 different scenarios that were established in collaboration with the Washington Department of Fish and Wildlife to examine the recovery status of wolves given different management actions (translocation, harvest, lethal removal) and other factors (out-of-state immigration, disease). For each scenario, we predicted wolf population dynamics from 2021-2070 and quantified metrics related to wolf population status, including the probability of meeting recovery criteria. Our approach allowed us to propagate both demographic and structural uncertainty into our 50-year forecasts of wolf population dynamics. Capturing this uncertainty is an important step when forecasting how management actions may influence management outcomes in order to assist managers making decisions under uncertainty (Runge and Converse 2020).

## METHODS

### Study area

Washington State is located on the northwestern coast of the United States, bordered by British Columbia, Canada to the north, Idaho to the east, and Oregon to the south. There are three recovery regions in Washington’s Wolf Conservation and Management Plan: Eastern Washington, the Northern Cascades, and the Southern Cascades and Northwest Coast (Wiles et al. 2011). There are currently (as of December 2022) wolf packs in the former two but not the latter recovery region (Figure 1).

**Figure 1.**
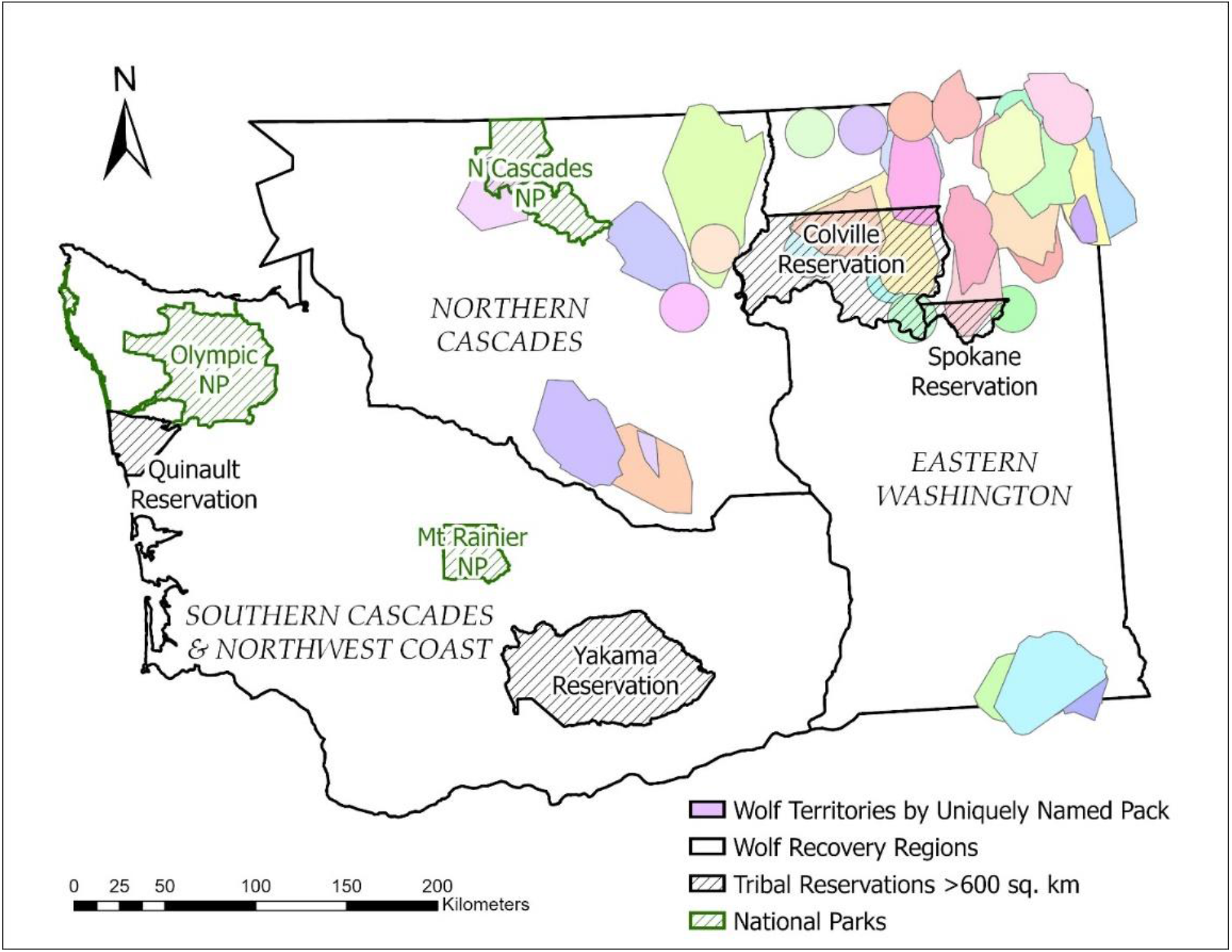
Thirty-eight uniquely named packs and the three wolf recovery regions in Washington State during the data collection period (2009-2020). These 38 named packs correspond to 34 pack territories occupied during annual surveys conducted in winter each year. Some pack names changed for the same territory due to the pack naturally dissolving or lethal removals due to livestock depredations. Packs represented by circles are those that were known to exist but for which individual movement data from Global Positioning System collars were not available.

### Projection model

Petracca et al. (2024) describes a spatially explicit projection model that combines (1) an integrated population model for demographic estimation and (2) an individual-based model for projecting movement and recolonization of wolves across a landscape of hypothetical pack territories in Washington. The habitat suitability of each hypothetical pack territory was determined based on a resource selection function. We used multiple data sources to estimate parameters of the model, including movement and demographic data from Global Positioning System (GPS) collars placed on 74 individual wolves by the Washington Department of Fish and Wildlife (WDFW) from 2009 to 2020, along with pack counts for these years (Petracca et al. 2024). The data represented 38 packs in Washington, where a “pack” is defined as two or more wolves traveling together in winter. Here, we used the same model structure along with empirical estimates of model parameters in our population projections. While Petracca et al. (2024) considers two different models for selection of new territories by dispersing wolves, here we used the “RSF Categorical” approach, in which wolves selected territories based on the quality of habitat available in all potential new territories at a given distance from the origin territory.

We used estimates of reproduction and movement rates from the estimation model in Petracca et al. (2024), while the projected survival in each six-month period *t* was sampled from a normal hyper-distribution with hyper-parameters estimated on a logit scale, and then transformed to the probability scale:

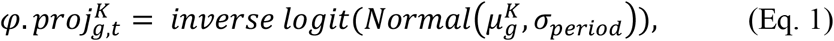

Where 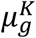 was the logit-scale mean six-month survival probability of animals in age group *g* (juvenile, adult) and movement state *K* (resident, mover), and *σ*_*period*_ was the logit-scale standard deviation for the random effect of six-month period, both estimated over the data collection period. Baseline parameter values for removal rate, out-of-state immigration rate, reproduction rate, movement rate, and survival, estimated in Petracca et al. (2024), are provided in Table 1.

**Table 1.** Baseline mean and 95% credible intervals for out-of-state immigration rate, reproduction rate, movement rate, and survival, as estimated in Petracca et al. (2024). Lethal removal rate was calculated directly from state agency records.

| Description | Mean (95% Credible Interval, if applicable) |
| --- | --- |
| Out of state immigration rate, i.e., # of new immigrants per occupied territory in 6-mo period | 0.11 (0.01 - 0.28) |
| Prob of a territory with 2+ wolves having 0 6-mo-olds | 0.50 (0.38 - 0.62) |
| ...1 6-mo-olds | 0.07 (0.02 - 0.16) |
| ...2 6-mo-olds | 0.14 (0.06 - 0.25) |
| ...3 6-mo-olds | 0.17 (0.09 - 0.26) |
| ...4 6-mo-olds | 0.08 (0.03 - 0.15) |
| ...5 6-mo-olds | 0.02 (0.00 - 0.06) |
| ...6 6-mo-olds | 0.02 (0.00 - 0.05) |
| Prob of initiating movement for residents 12-24 mos | 0.38 (0.25 - 0.52) |
| Prob of initiating movement for residents $\geq 24$ mos | 0.06 (0.03 - 0.10) |
| Prob of continuing movement for movers 12-24 mos | 0.33 (0.26 - 0.42) |
| Prob of continuing movement for movers $\geq 24$ mos | 0.43 (0.35 - 0.51) |
| Prob of mover wolf staying within WA State | 0.75 (0.64 - 0.85) |
| 6-mo survival of residents 6-24-mo-olds (see Eq 1) | 0.98 (0.91 - 1.00) |
| 6-mo survival of residents $\geq 24$ mos (see Eq 1) | 0.94 (0.89 - 0.98) |
| 6-mo survival of movers 12-24 mo-old (see Eq 1) | 0.84 (0.46 - 1.00) |
| 6-mo survival of movers $\geq 24$ mos (see Eq 1) | 0.98 (0.92 - 1.00) |
| Annual lethal removal rate | 0.04 |

### Population viability analysis and management scenarios

For the projection period (2021-2070), we applied the baseline scenario as in Petracca et al. (2024) and added 11 additional scenarios. All scenarios began with estimated wolf abundance from December 2020 (median of 172, 95% prediction interval [154-191]), the last year for which we had data. Scenarios were identified in collaboration with the Washington Department of Fish and Wildlife (Table 2). For each scenario, we predicted spatially referenced abundance and summarized these predictions to calculate population growth rate, the probability of meeting recovery objectives, quasi-extinction probability, and extinction probability.

**Table 2.**
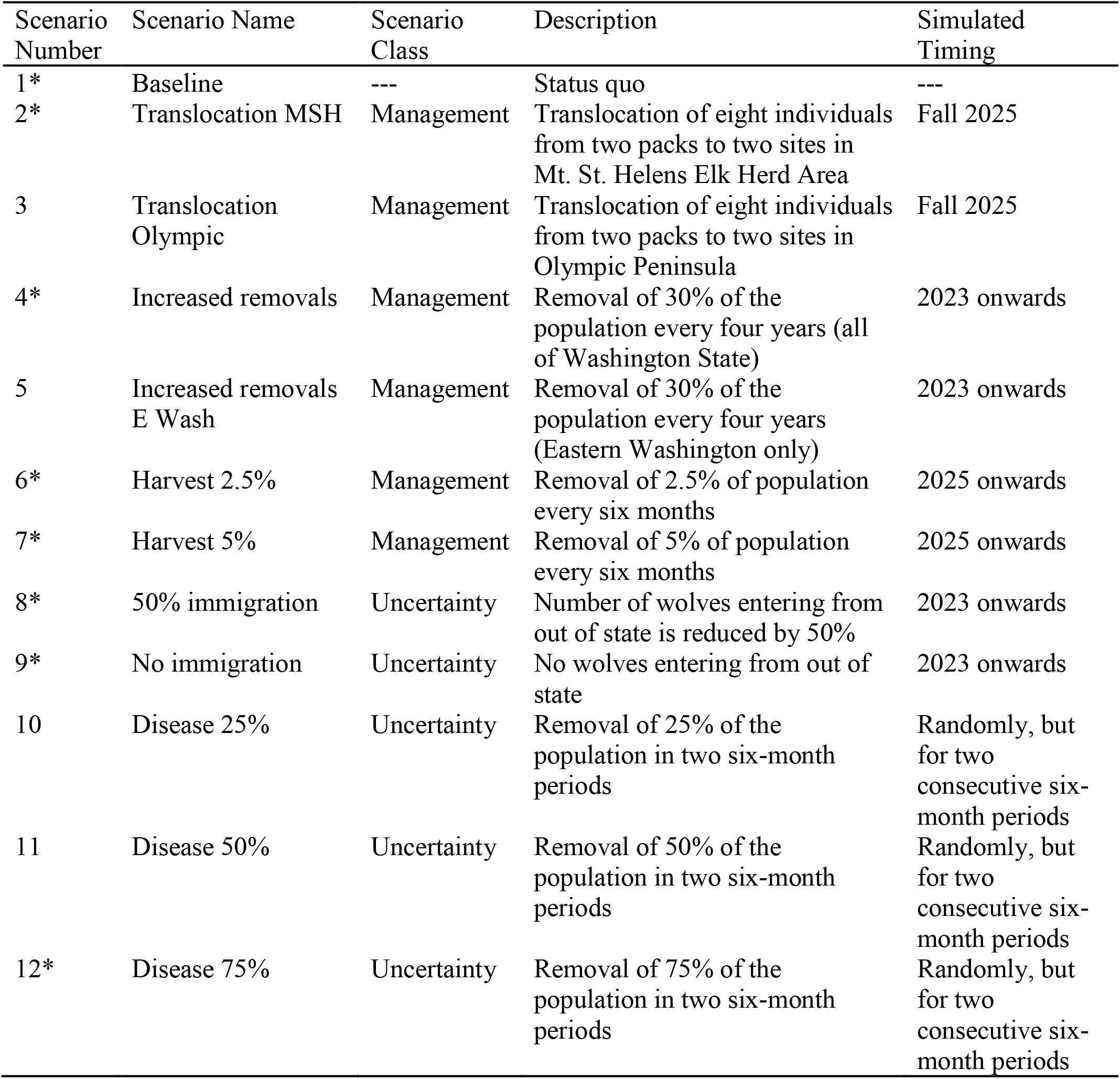
Summary of 12 scenarios used to project the gray wolf population of Washington State from 2021-2070 using the spatially explicit projection model described in Petracca et al. (2024). Scenario numbers with (*) were the focus of the main text, with all other scenario results presented in the Appendix. Scenarios were either a target for management (“Management”) or a consideration of system uncertainty (“Uncertainty”).

### Baseline

In scenario 1 (“Baseline”) we simulated all relevant factors, as described below, at levels observed in the data collection period (2009-2020). We assumed maintenance of lethal removals (wolf removals by the state agency in response to livestock depredations [WDFW 2020]) at the mean annual rate from 2009-2020 (3.5%), with removals applied to the Eastern Washington recovery area only. Eastern Washington is where Washington currently has the most wolves, as well as private lands, and where lethal removals presently occur. For a given number of annual removals, a pack was chosen at random and all individuals from that pack were removed; any remaining removals were removed at random from another randomly selected pack. We also maintained the same number of wolves immigrating into Washington from out of state as was estimated across the data period. We assumed no harvest or translocations. For all other scenarios, conditions were left as in the baseline except for changes as noted.

### Translocation

In scenarios 2 and 3 (“Translocation”), we simulated moving eight wolves (two groups of four, each group having two 30+-month-old adults and two 6-month-old pups) from two randomly selected packs in Eastern Washington to the South Cascades and Northwest Coast recovery region. In scenario 2, we simulated moving the eight wolves to the Mt St Helens Elk Herd Area in autumn of year 5 of the simulation (i.e., in 2025), specifically to the two hypothetical pack territories anywhere in the South Cascades and Northwest Cost recovery region with the highest estimated suitability based on the resource selection function presented in Petracca et al. (2024). In scenario 3, we simulated moving eight wolves to the Olympic Peninsula in autumn of year 5 to the two hypothetical pack territories on the Peninsula with the highest estimated suitability.

### Increased removals

In scenarios 4 and 5 (“Increased removals”), we simulated an increased number of lethal management removals such that 30% of the wolf population would be removed every four years, corresponding to an annual removal rate of 8.5%. In Scenario 4, removals applied to all wolves in Washington. In Scenario 5, removals applied to wolves in the Eastern Washington recovery region only, as removals are currently restricted to that recovery region. The increased removal rate was applied starting in year 3 of the simulation (i.e., in 2023), and continued throughout the 50-year simulation.

### Harvest

In scenarios 6 and 7 (“Harvest”), we simulated the introduction of legal harvest such that 2.5% or 5% of the entire population was removed every six months. We simulated this mortality as fully additive, such that harvest mortality occurs independently of other forms of mortality (Mills 2013). These scenarios directly manipulated survival of individuals in both age groups (*g*, juvenile or adult) and both movement states *(K*, resident or mover) in six-month time period 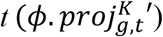 such that

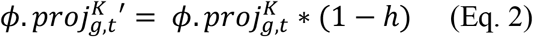

where harvest level *h* was either 0.025 or 0.05, and 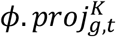 is the survival rate without harvest, as described in Eq. 1. Scenario 6 simulated 2.5% removal every six months and Scenario 7 simulated 5% removal every six months. We began harvest in year 5 of the simulation (i.e., in 2025).

### Immigration

In scenarios 8 and 9, we varied the immigration rate into Washington. In scenario 8 (“50% immigration”), we simulated a 50% reduction in immigration of wolves into Washington. We multiplied the parameter *λ*_*immig*_ (number of immigrants per six-month-period per pack; see Petracca et al. [2024]) by 0.5 for all samples. This process was simulated to start in year 3 of the simulation (i.e., in 2023) and continue for the length of the simulation. In scenario 9 (“No immigration”), we simulated complete elimination of immigration of wolves into Washington. In scenario 9, we set the parameter *λ*_*immig*_ to 0 for all samples, also starting in year 3 of the simulation (i.e., in 2023).

### Disease

In scenarios 10 through 12 (“Disease”), we simulated an increase in mortality due to disease. We removed 25%, 50%, or 75% of the entire population. Again, we assumed this mortality to be fully additive. Similar to harvest, these scenarios directly manipulated survival of individuals in state *K* of age grouping *g* in six-month time period 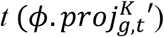 such that

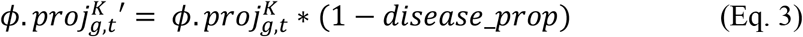

where 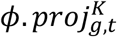 is the survival rate without disease, as described in Eq. 1. Proportion of the population removed by disease *disease_prop* was 0.25 (Scenario 10), 0.50 (Scenario 11), or 0.75 (Scenario 12). Disease was estimated to occur at a random time between year 3 (i.e., June 2023) and the end of the simulation, and to persist for two consecutive six-month periods.

### Implementation

The projection model was run in R v.4.2.0 (R Core Team 2022) for 50 years, using 500 samples from the posterior distributions of the model parameters (Table 1) with 100 stochastic simulations per sample. Metrics for each simulation included total number of wolves, geometric mean of population growth rate, probability of reaching the recovery threshold across all years (2021-2070), probability of dropping below the quasi-extinction threshold across all years (2021-2070), and the probability of extinction. Quasi-extinction was defined as having <92 adult wolves in the state and <24 adult wolves in each of three recovery regions (Wiles et al. 2011). Extinction was defined as having zero wolves in Washington State in year 50 (i.e., 2070).

## RESULTS

Due to similarities in projections among scenario classes, we present results from eight scenarios here (Table 2): Baseline, Translocation MSH, Increased removals, Harvest 2.5%, Harvest 5%, 50% immigration, No immigration, and Disease 75%. Please see Appendix S1 for results from all 12 modeled scenarios.

Median probability of recovery (i.e., four breeding pairs in each recovery region, with three additional breeding pairs anywhere in the state) across all years (2021-2070) was above 50% for the Baseline (0.66, 95% PI 0.02-0.88) and Translocation MSH (0.67, 0.02-0.88) scenarios, and below 50% for the 50% immigration (0.48, 0.00-0.85), Disease 75% (0.38, 0.01-0.70), Harvest 2.5% (0.31, 0.00-0.85), No immigration (0.12, 0.00-0.81), Increased removals (0.12, 0.00-0.72), and Harvest 5% (0.07, 0.00-0.64) scenarios (Figure 2).

**Figure 2.**
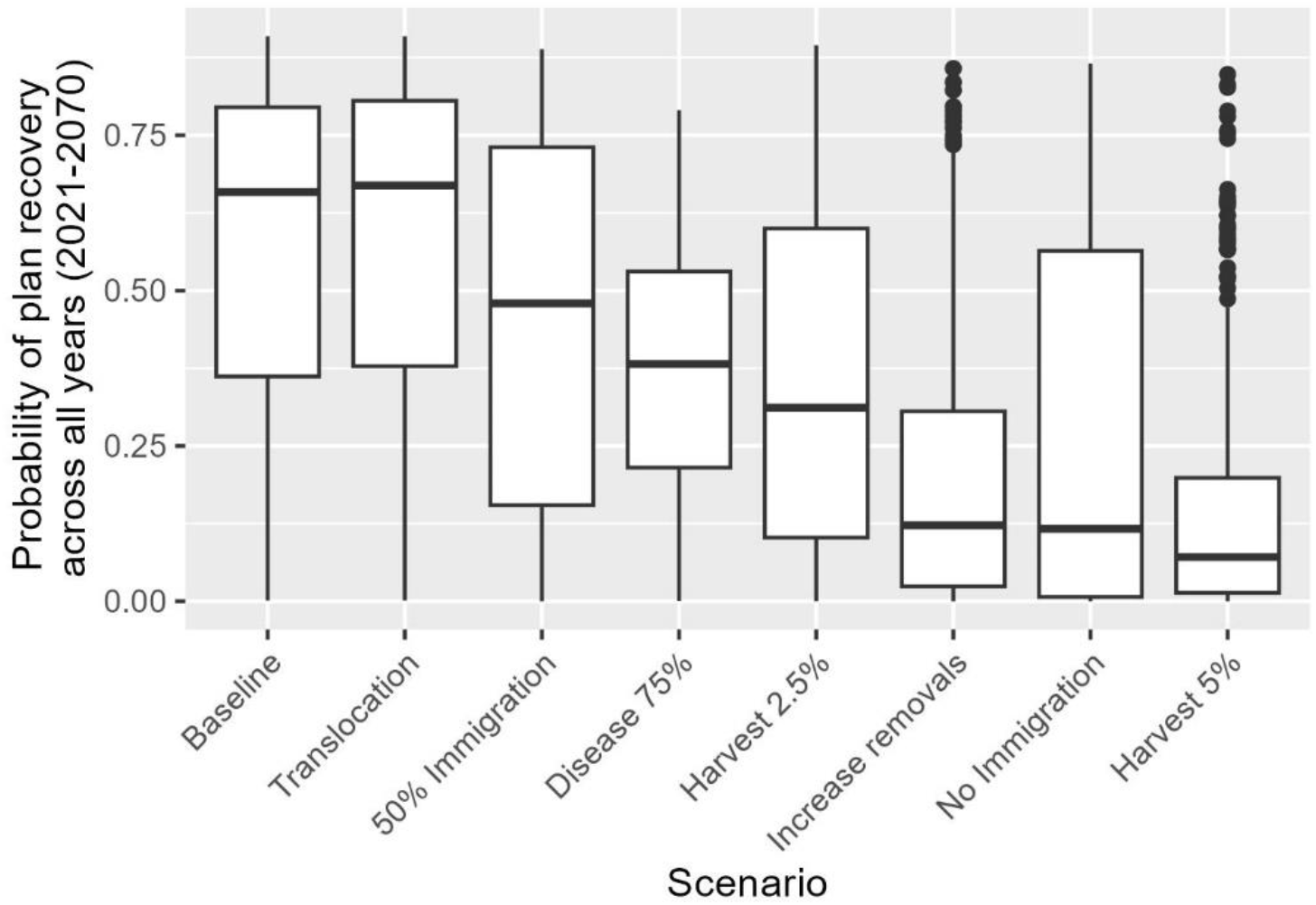
Probability of meeting plan recovery across all years (2021-2070) for seven scenarios related to management and system uncertainty. Plan recovery is considered having four breeding pairs in each recovery region, with three additional breeding pairs anywhere in the state. The center line represents the median and boxes represent the 50% prediction interval. The points represent individual data points (of 50,000 samples) that are outliers.

Median probability of recovery increased from the baseline for all scenarios but Harvest 5% (Figure 3). Probability of recovery was >50% by the year 2040 for the Baseline and Translocation scenarios, and by 2060 for 50% immigration; the other five scenarios (No immigration, Increased removals, Harvest 2.5%, Harvest 5%, Disease 75%) failed to reach the >50% probability of recovery threshold by 2070.

**Figure 3.**
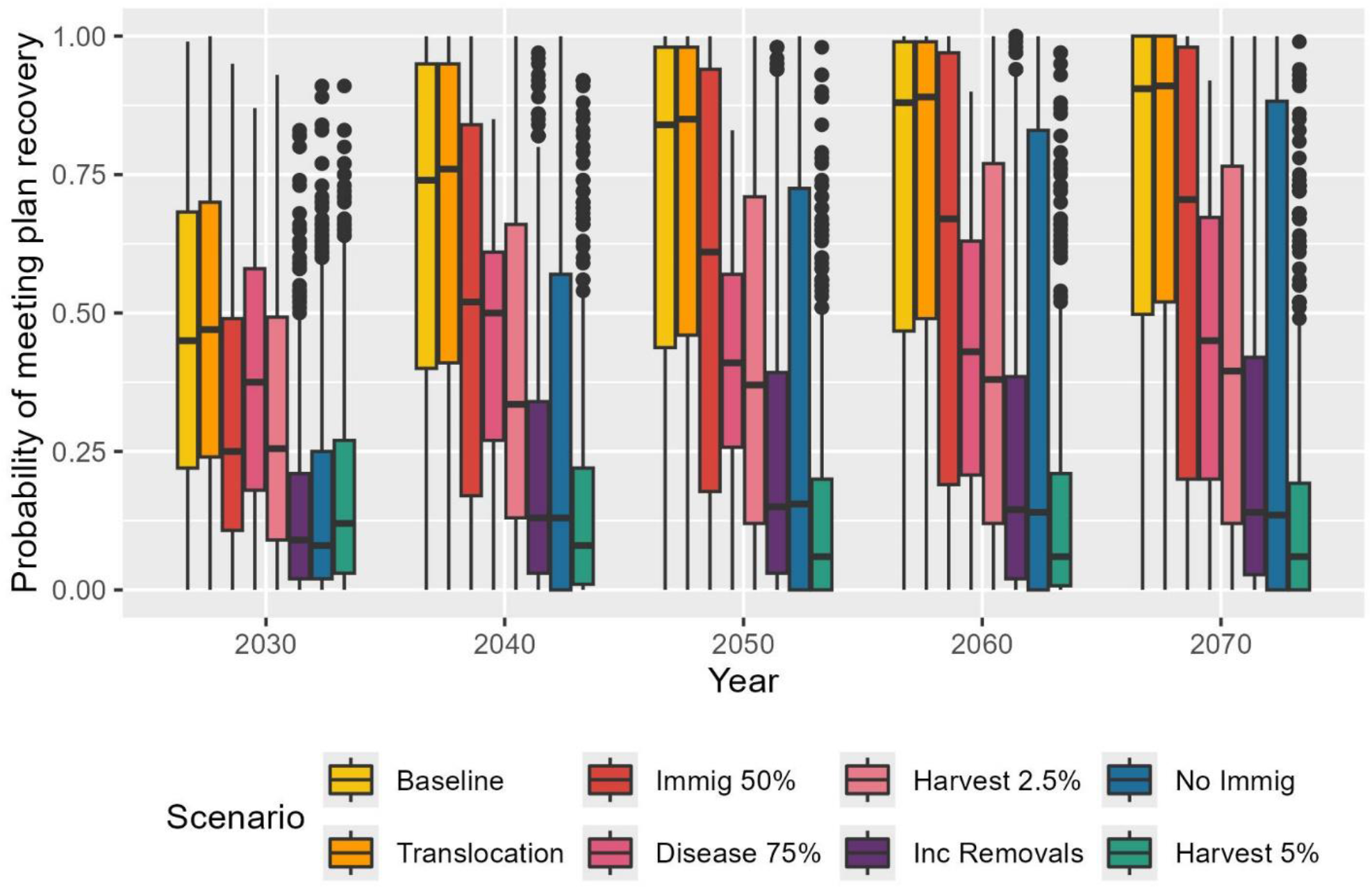
Probability of meeting plan recovery at various time points over the period 2021-2070 for seven scenarios related to management and system uncertainty. Plan recovery is defined as having 4 breeding pairs in each recovery region, with 3 additional breeding pairs anywhere in the state. The center line represents the median and boxes represent the 50% prediction interval. The points represent individual data points (of 50,000 samples) that are outliers.

The geometric mean of lambda was above 1 for all scenarios but No immigration (0.99; 95% PI 0.83-1.04; Figure 4). Geometric mean of lambda was 1.03 (95% PI 1.00-1.05) for two scenarios: Baseline and Translocation. Geometric mean of lambda was 1.05 (1.01-1.07) for Disease 75%, 1.02 (1.00-1.05) for 50% Immigration, 1.02 (0.99-1.04) for Harvest 2.5%, 1.01 (0.97-1.04) for Increased removals, and 1.01 (0.97-1.04) for Harvest 5%.

**Figure 4.**
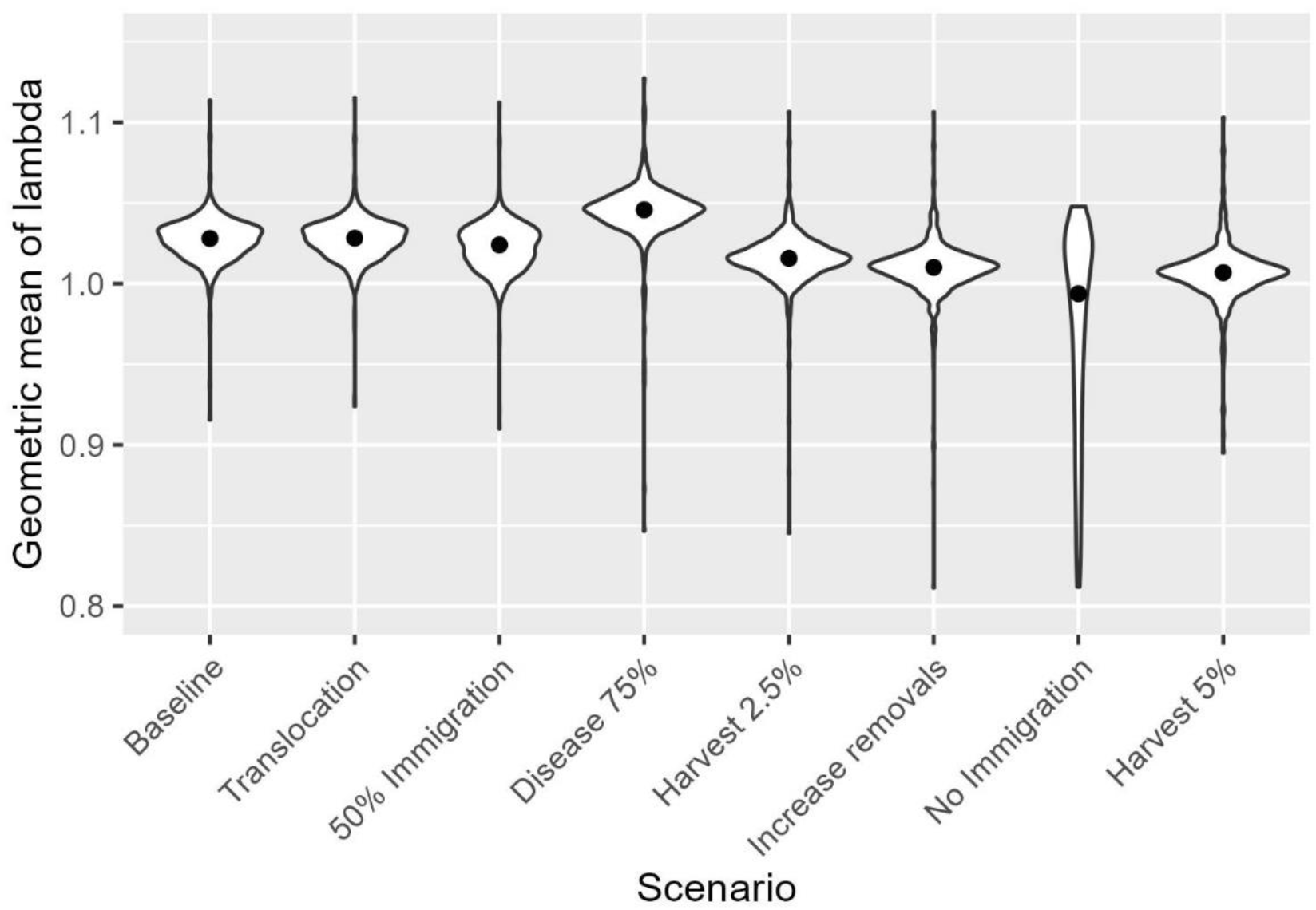
Projected geometric mean of population growth, *λ*, for gray wolves (*Canis lupus*) population in Washington State, USA, over the time period 2021-2070, for eight scenarios related to management and system uncertainty.

Median probability of extinction (i.e., zero wolves in 2070) was zero across all scenarios but No immigration (0.05, 95% PI 0.00-1.00) (Appendix S1). Predicted probabilities of quasi-extinction (i.e., <92 adult wolves in the state and <24 adult wolves in each recovery region in any year) across all years (2021-2070) were low (<20%) across all scenarios. Median probability of quasi-extinction was 0.18 (95% PI 0.00 – 0.83) for No immigration, 0.09 (95% PI 0.03-0.54) for Disease 75%, 0.04 (0.00-0.73) for Harvest 5%, 0.02 (0.00-0.68) for Increased removals, and 0.01 (0.00-0.59) for Harvest 2.5%. All other scenarios had a median probability of quasi extinction of 0, with varying prediction intervals (0.00 [95% PI 0.00-0.39] for Baseline, 0.00 [0.00 -0.37] for Translocation MSH, and 0.00 [0.00-0.57] for 50% immigration (Figure 5).

**Figure 5.**
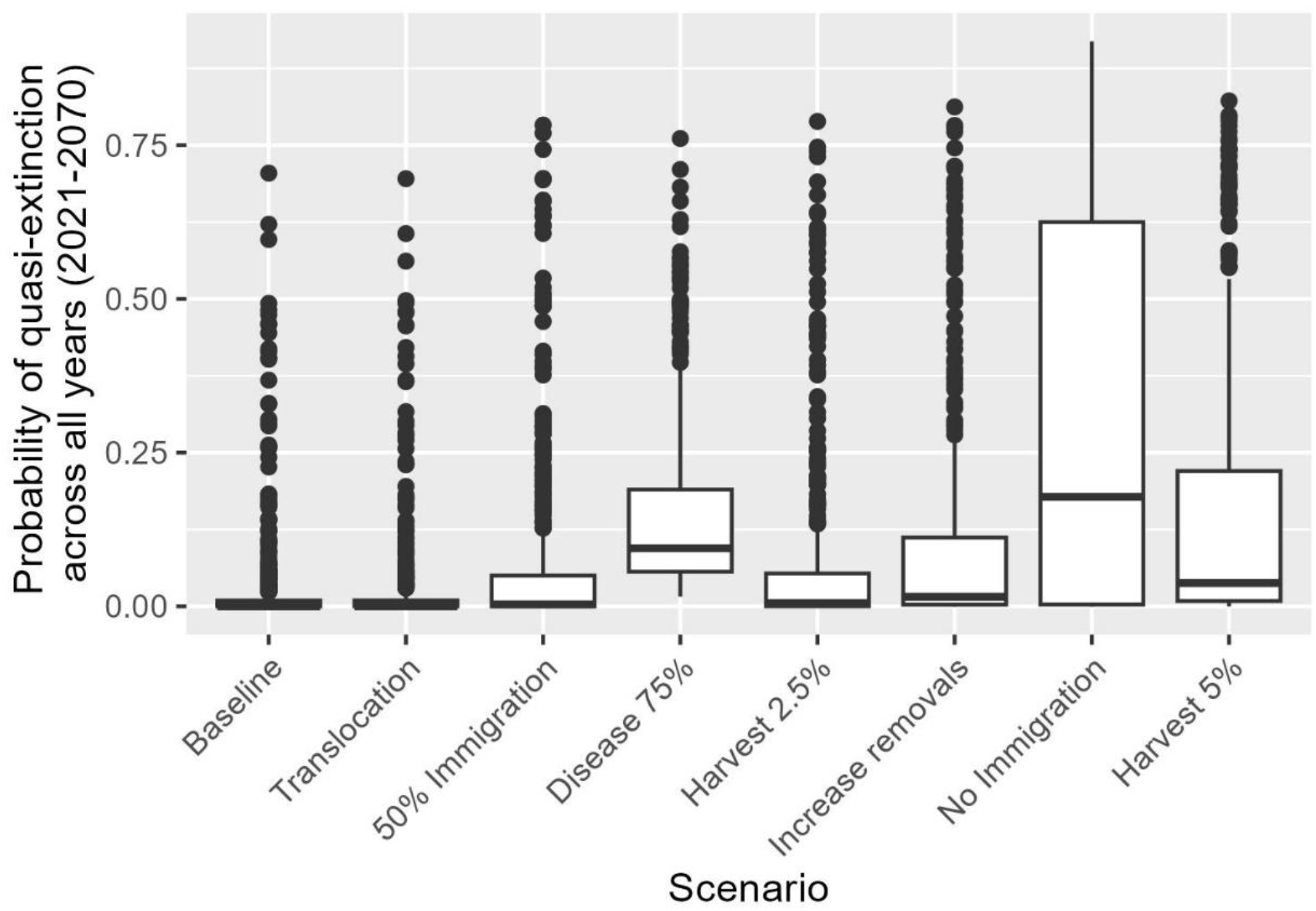
Probability of quasi-extinction across all years (2021-2070) for seven scenarios related to management and system uncertainty. Quasi-extinction occurs when there are <92 adult wolves in the state and <24 adult wolves in each recovery region. The center line represents the median and boxes represent the 50% prediction interval. The points represent individual data points (of 50,000 samples) that are outliers.

All scenarios started with a median wolf abundance of 172 (95% prediction interval [PI] 154-191) in the year 2020. Predicted wolf abundance in 2070 was 469 (50-1261) for Baseline, 475 (95% PI 51-1271) for Translocation, 357 (25-1204) for 50% immigration, 237 (18-714) for Harvest 2.5%, 226 (10-904) for Disease 75%, 177 (8-462) for Increased removals, 140 (5-362) for Harvest 5%, and 99 (0-1133) for No immigration (Figure 6).

**Figure 6.**
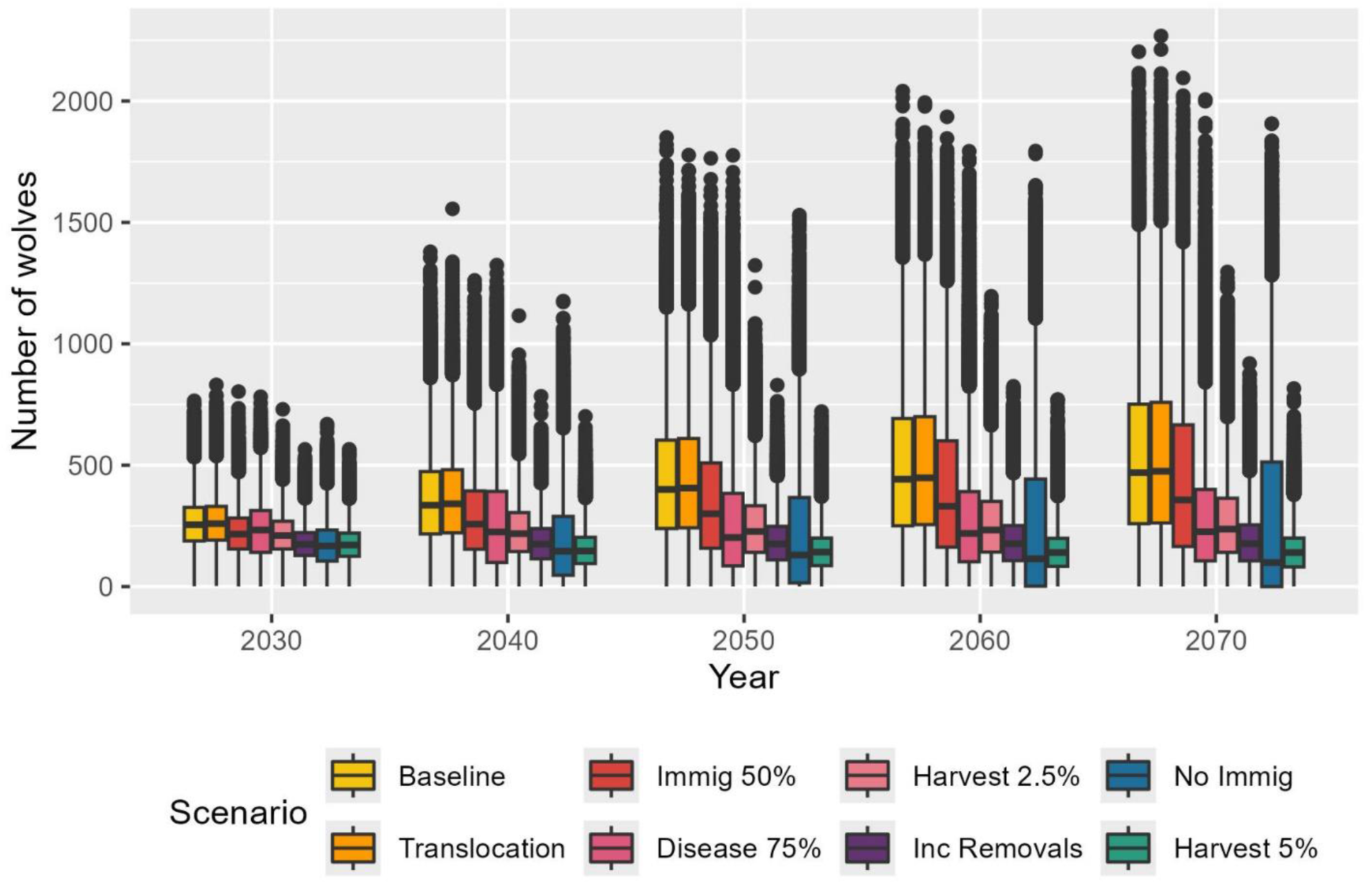
Estimated number of wolves in Washington State, USA, for seven scenarios related to management and system uncertainty. Black line indicates the median and boxes represent the 50% prediction interval. The points represent individual data points (from 50,000 samples) that are outliers.

## DISCUSSION

Predicting the trajectory of recovering species is notoriously challenging and is made more so by the wide variety of factors – both within and outside the control of managers – that may influence recovering populations (Coulson et al. 2001, Ellner et al. 2002, McCarthy et al. 2003). Population projections are a powerful tool for exploring the effects of uncertainties on species recovery and can inform the decisions of managers tasked with recovering species and determining their legal status (Morris and Doak 2002). Using an existing population projection framework for wolves in Washington State, we projected the population forward under 12 scenarios relating to management actions (lethal removals, translocation, harvest) and sources of structural uncertainty (out-of-state immigration, disease) and found that recovery criteria for wolves in Washington were likely to be met across the next 50 years under three of eight scenarios. In developing scenarios, we included a wide range of potential factors that could impact wolf population trajectories and found that the population’s long-term growth rate was robust to possible disease outbreaks, harvest, and lethal removals. These results are consistent with the success of recolonizing wolf populations across North American and Europe including in the upper Midwestern United States (Mech 1995), northern Rocky Mountains of the United States (Smith et al. 2010, Jimenez et al. 2017), Scandinavia (Vilà et al. 2003, Wabakken et al. 2011), and mainland Europe (Chapron et al. 2014). We note that our model assumed vital and movement rates in the projection period (2021-2070) to be drawn from the same distribution as those observed during our data collection period (2009-2020), unless those rates were specifically affected by the management strategy.

While population growth rates across scenarios were indicative of a stable or growing population, some scenarios had a relatively low probability of meeting recovery objectives. The three scenarios that resulted in the lowest probabilities of meeting recovery criteria across all years (2021-2070) were the Harvest 5% (7% recovery probability), Increased removals (12%), and the No immigration (12%) scenarios. The harvest and removal scenarios likely had the lowest probability of meeting recovery goals due to offtake (5% of the population removed every six months for harvest, 30% of the population removed every four years for lethal removals) being too high to sustain the population (see Creel et al. [2015] and rebuttal by Mitchell et al. [2016] for policy considerations regarding wolf offtake). The two strategies that we simulated specified a fixed harvest and lethal removal rate, respectively, across all years and regions; a more likely management approach would involve adaptation of these rates if the population was estimated to be declining (Andrén et al. 2020). In addition, some mortality may be compensatory (Murray et al. 2010), which we did not attempt to account for given the challenges of parameterizing a mortality process that is compensatory. In other scenarios, we assumed that harvest was 0%, as it is not currently permitted in Washington outside tribal areas (Washington Department of Fish and Wildlife, Confederated Tribes of the Colville Reservation, Spokane Tribe of Indians, USDA-APHIS Wildlife Services, and U.S. Fish and Wildlife Service 2022). Legal harvest by tribal members has occurred in Washington (mean annual harvest = 4.83 [SD = 2.14] wolves statewide from 2015-2020); these mortalities were accounted for in our background mortality rate.

Immigration is an important source of new individuals in a recovering population (Altwegg et al. 2014, Lieury et al. 2016, Grauer et al. 2019), and has been critical to the recolonization of wolves in Washington State (Maletzke et al. 2016). The recent increase in wolf harvest in neighboring Idaho (Idaho State Legislature 2021), Montana (MFWP 2022), and British Columbia (Government of Canada 2020) has led to an increase in questions regarding whether current immigration levels will remain. While our No immigration scenario was extreme, enforcing a complete cessation of out-of-state immigration rather than a reduction, it does underscore the importance of out-of-state immigration to sustaining the population within Washington (though see Bassing et al. (2020) for how immigration alone cannot compensate for high anthropogenic mortality). Under the 50% immigration scenario, by contrast, which is arguably more realistic and still enforced a large decline in immigration, we predicted a 71% median probability that the population would meet recovery criteria in 2070. These scenarios show not only the importance of out-of-state immigration to the Washington wolf population, but also that uncertainty around immigration makes prediction of recovery challenging.

The other two scenarios with <50% probability of meeting recovery goals across all years (2021-2070) were those involving short-term (i.e., Disease 75%) or more modest (i.e., Harvest 2.5%) reductions in survival. In our disease scenarios, we considered up to 75% reduction in survival in two random consecutive six-month periods. We applied this increased mortality to all age classes; however, in some cases, diseases are more likely to impact one age class than another. For example, canine parvovirus has been shown to reduce pup survival by 40-60% while not directly causing mortality of older age classes (Mech et al. 2008); canine distemper has also disproportionately impacted pup survival (Almberg et al. 2009). As younger individuals are more likely to disperse, the results of such outbreaks may also reduce the recolonization rate of wolves (Mech et al. 2008). Consideration of how different diseases may impact the population is important for understanding not only demographic rates, but also dispersal and recolonization rates.

One commonly used management tool to help small or extirpated populations recolonize an area is conservation translocation (Seddon et al. 2007, Seddon and Armstrong 2016). Our translocation scenarios, which involved relatively modest numbers of wolves in translocated cohorts, were not predicted to discernibly change progress toward recovery. This lack of measurable impact occurred despite translocated wolves being assumed to have survival rates drawn from the same distribution as resident wolves elsewhere in the state, though rates for translocated individuals may be lower in practice (Bradley et al. 2005). Given that the population was predicted to expand under the baseline scenario, the relatively small number of translocated individuals was unlikely to accelerate recovery. Larger translocation cohorts may be more effective in accelerating recovery by increasing the establishment of wolves in the Southern Cascades and Northwest Coast recovery region, but translocation of wolves is a politically and logistically challenging undertaking.

While wolves continue to expand in range, conflicts between people and wolves are likely to follow suit (Mech 1995, Breitenmoser 1998). This is particularly true in Washington, where the urban-wildland gradient is changing quickly and wolves are settling in areas with more livestock grazing (Hanley et al. 2018). Human-wolf conflicts arise due to issues such as livestock depredation, competition for hunted game species, and the perceived threat of potential direct attacks on humans (Treves and Karanth 2003, Rodríguez-Recio et al. 2022). Currently, the Washington Department of Fish and Wildlife uses lethal removals in response to livestock depredation, and we considered an increase in lethal removals in our scenarios to account for this potential increase in human-wildlife conflict. While lethal removals currently only occur in the Eastern Washington recovery region, it is likely that removals may become more common in the Northern Cascades given its high predicted likelihood of wolf-livestock interactions as the wolf population expands (Hanley et al. 2018). It is also important to consider that lethal removals may not increase linearly with wolf population size as we modeled here, and that use of non-lethal deterrents to prevent livestock loss – such as flagging, aversive stimuli (e.g., shock collars, less-than-lethal munitions), and disruptive stimuli (e.g., light and siren devices) – may provide deterrents to livestock depredation (Bangs and Shivik 2001, Shivik et al. 2003, Bangs et al. 2006).

Accounting for uncertainty is necessary to provide robust predictions of population trajectories (Morris and Doak 2002, McGowan et al. 2011). Here, we accounted for uncertainty related to both lack of knowledge or information about the system (i.e., epistemic uncertainty) and uncertainty related to the inherent randomness of the system (i.e., aleatory uncertainty) (Regan et al. 2002). We captured epistemic uncertainty by exploring multiple structural uncertainties related to immigration and disease. We captured aleatory uncertainty by accounting for (1) parametric uncertainty via the posterior distributions of model parameters as estimated in Petracca et al. (2024), (2) environmental stochasticity by generating future survival rates from hyperparameters and allowing disease to occur stochastically (see Eq. 1), and (3) demographic stochasticity through the stochastic structure of the model described in Petracca et al. (2024). While the result is substantial uncertainty about population trajectories, this uncertainty accurately characterizes the reality faced by managers (Morris and Doak 2002). Methods for decision making in the face of risk (e.g., Runge and Converse 2020) are available to help decision makers grapple with this uncertainty.

We used a spatially explicit projection model to predict the spatiotemporal dynamics of a recolonizing carnivore in Washington State, USA, conditional on a variety of factors both within and outside manager’s control. Our results suggest that a recolonizing carnivore with typically high reproductive rates and dispersal ability is robust to some management strategies and sources of uncertainty, but is unlikely to recover within 50 years in scenarios of high harvest, high lethal management removals, and cessation of immigration. This exercise underscored the important role played by immigration and highlighted the potential risks of high harvest and lethal management removals for sustaining a recolonizing population, a lesson that can be applied broadly across threatened taxa.

## ACKNOWLEDGMENTS

We thank the Washington Department of Fish and Wildlife (WDFW) for providing funding for this work, and the Washington Cooperative Fish and Wildlife Research Unit for facilitating the funding. We deeply appreciate the insights and advice of WDFW scientists and managers, particularly D Martorello, T Roussin, J Smith, and G Spence, and we thank the members of the Washington Fish and Wildlife Commission, particularly the Wolf Committee, for challenging us to continually improve this product. Members of the Quantitative Conservation Lab and the Quantitative Ecology Lab at the University of Washington provided advice and support throughout. Any use of trade, firm, or product names is for descriptive purposes only and does not imply endorsement by the U.S. Government.

## REFERENCES

Abrahms, B., K. Rafiq, N. R. Jordan, and J. W. McNutt. 2022. Long-term, climate-driven phenological shift in a tropical large carnivore. Proceedings of the National Academy of Sciences 119:e2121667119.

Almberg, E. S., L. D. Mech, D. W. Smith, J. W. Sheldon, and R. L. Crabtree. 2009. A Serological Survey of Infectious Disease in Yellowstone National Park’s Canid Community. PLOS ONE 4:e7042.

Altwegg, R., A. Jenkins, and F. Abadi. 2014. Nestboxes and immigration drive the growth of an urban Peregrine Falcon Falco peregrinus population. Ibis 156:107–115.

Andrén, H., N. T. Hobbs, M. Aronsson, H. Brøseth, G. Chapron, J. D. C. Linnell, J. Odden, J. Persson, and E. B. Nilsen. 2020. Harvest models of small populations of a large carnivore using Bayesian forecasting. Ecological Applications 30:e02063.

Armstrong, D. P., and P. J. Seddon. 2008. Directions in reintroduction biology. Trends in Ecology & Evolution 23:20–25.

Bangs, E. E., and S. H. Fritts. 1996. Reintroducing the gray wolf to central Indaho and Yellowstone National Park. Wildlife Society Bulletin 24:402–413.

Bangs, E. E., and J. A. Shivik. 2001. Managing wolf conflict with livestock in the northwestern United States. Carnivore Damage Prevention News 3:2–5.

Bangs, E., M. Jimenez, C. Niemeyer, J. Fontaine, M. Collinge, R. Krsichke, L. Handegard, J. A. Shivik, C. Sime, S. Nadeau, C. Mack, D. W. Smith, V. Asher, and S. Stone. 2006. Non-Lethal and Lethal Tools to Manage Wolf-Livestock Conflict in the Northwestern United States. Proceedings of the Vertebrate Pest Conference 22.

Bassing, S. B., D. E. Ausband, M. S. Mitchell, M. K. Schwartz, J. J. Nowak, G. C. Hale, and L. P. Waits. 2020. Immigration does not offset harvest mortality in groups of a cooperatively breeding carnivore. Animal Conservation 23:750–761.

Bradley, E. H., D. H. Pletscher, E. E. Bangs, K. E. Kunkel, D. W. Smith, C. M. Mack, T. J. Meier, J. A. Fontaine, C. C. Niemeyer, and M. D. Jimenez. 2005. Evaluating Wolf Translocation as a Nonlethal Method to Reduce Livestock Conflicts in the Northwestern United States. Conservation Biology 19:1498–1508.

Breitenmoser, U. 1998. Large predators in the Alps: The fall and rise of man’s competitors. Biological Conservation 83:279–289.

Chapron, G., P. Kaczensky, J. D. C. Linnell, M. von Arx, D. Huber, H. Andren, J. Vicente Lopez-Bao, M. Adamec, F. Alvares, O. Anders, L. Balciauskas, V. Balys, P. Bedo, F. Bego, J. Carlos Blanco, U. Breitenmoser, H. Broseth, L. Bufka, R. Bunikyte, P. Ciucci, A. Dutsov, T. Engleder, C. Fuxjaeger, C. Groff, K. Holmala, B. Hoxha, Y. Iliopoulos, O. Ionescu, J. Jeremic, K. Jerina, G. Kluth, F. Knauer, I. Kojola, I. Kos, M. Krofel, J. Kubala, S. Kunovac, J. Kusak, M. Kutal, O. Liberg, A. Majic, P. Maennil, R. Manz, E. Marboutin, F. Marucco, D. Melovski, K. Mersini, Y. Mertzanis, R. W. Myslajek, S. Nowak, J. Odden, J. Ozolins, G. Palomero, M. Paunovic, J. Persson, H. Potocnik, P.-Y. Quenette, G. Rauer, I. Reinhardt, R. Rigg, A. Ryser, V. Salvatori, T. Skrbinsek, A. Stojanov, J. E. Swenson, L. Szemethy, A. Trajce, E. Tsingarska-Sedefcheva, M. Vana, R. Veeroja, P. Wabakken, M. Woefl, S. Woelfl, F. Zimmermann, D. Zlatanova, and L. Boitani. 2014. Recovery of large carnivores in Europe’s modern human-dominated landscapes. Science 346:1517–1519.

Coulson, T., G. M. Mace, E. Hudson, and H. Possingham. 2001. The use and abuse of population viability analysis. Trends in Ecology & Evolution 16:219–221.

Creel, S., M. Becker, D. Christianson, E. Dröge, N. Hammerschlag, M. W. Hayward, U. Karanth, A. Loveridge, D. W. Macdonald, W. Matandiko, J. M’soka, D. Murray, E. Rosenblatt, and P. Schuette. 2015. Questionable policy for large carnivore hunting. Science 350:1473–1475.

DeCesare, Nicholas. J., S. M. Wilson, E. H. Bradley, J. A. Gude, R. M. Inman, N. J. Lance, K. Laudon, A. A. Nelson, M. S. Ross, and T. D. Smucker. 2018. Wolf-livestock conflict and the effects of wolf management. The Journal of Wildlife Management 82:711–722.

Ditmer, M. A., R. M. Niemiec, G. Wittemyer, and K. R. Crooks. 2022. Socio-ecological drivers of public conservation voting: Restoring gray wolves to Colorado, USA. Ecological Applications 32:e2532.

Elbroch, L. M., P. E. Lendrum, J. Newby, H. Quigley, and D. J. Thompson. 2015. Recolonizing wolves influence the realized niche of resident cougars. Zoological Studies 54:1–11.

Ellner, S. P., J. Fieberg, D. Ludwig, and C. Wilcox. 2002. Precision of Population Viability Analysis. Conservation Biology 16:258–261.

Geffen, E., M. J. Anderson, and R. K. Wayne. 2004. Climate and habitat barriers to dispersal in the highly mobile grey wolf. Molecular Ecology 13:2481–2490.

Gordon, C. H., A. C. Banyard, A. Hussein, M. K. Laurenson, J. R. Malcolm, J. Marino, F. Regassa, A.-M. E. Stewart, A. R. Fooks, and C. Sillero-Zubiri. 2015. Canine Distemper in Endangered Ethiopian Wolves. Emerging Infectious Diseases 21:824–832.

Government of Canada. 2020. Southern Mountain Caribou in British Columbia: bilateral conservation agreement between Canada and British Columbia. <https://www.canada.ca/en/environment-climate-change/services/species-risk-public-registry/conservation-agreements/southern-mountain-caribou-british-colombia-2020.html.> Accessed 23 March 2023.

Grauer, J. A., J. H. Gilbert, J. E. Woodford, D. Eklund, S. Anderson, and J. N. Pauli. 2019. Modest immigration can rescue a reintroduced carnivore population. The Journal of Wildlife Management 83:567–576.

Hanley, Z. L., H. S. Cooley, B. T. Maletzke, and R. B. Wielgus. 2018. Cattle depredation risk by gray wolves on grazing allotments in Washington. Global Ecology and Conservation 16:e00453.

Hayes, R. D., and A. S. Harestad. 2000. Demography of a recovering wolf population in the Yukon. Canadian Journal of Zoology-Revue Canadienne De Zoologie 78:36–48.

Idaho State Legislature. 2021. Senate Bill No. 1211. <https://legislature.idaho.gov/wp-content/uploads/sessioninfo/2021/legislation/S1211.pdf.> Accessed 23 March 2023.

Jimenez, M. D., E. E. Bangs, D. K. Boyd, D. W. Smith, S. A. Becker, D. E. Ausband, S. P. Woodruff, E. H. Bradley, J. Holyan, and K. Laudon. 2017. Wolf dispersal in the Rocky Mountains, Western United States: 1993–2008. The Journal of Wildlife Management 81:581–592.

Lawler, J. J., S. P. Campbell, A. D. Guerry, M. B. Kolozsvary, R. J. O’Connor, and L. C. N. Seward. 2002. The Scope and Treatment of Threats in Endangered Species Recovery Plans. Ecological Applications 12:663–667.

Lieury, N., A. Besnard, C. Ponchon, A. Ravayrol, and A. Millon. 2016. Geographically isolated but demographically connected: Immigration supports efficient conservation actions in the recovery of a range-margin population of the Bonelli’s eagle in France. Biological Conservation 195:272–278.

Maletzke, B. T., R. B. Wielgus, D. J. Pierce, D. A. Martorello, and D. W. Stinson. 2016. A meta-population model to predict occurrence and recovery of wolves. The Journal of Wildlife Management 80:368–376.

McCarthy, M. A., S. J. Andelman, and H. P. Possingham. 2003. Reliability of Relative Predictions in Population Viability Analysis. Conservation Biology 17:982–989.

McGowan, C. P., M. C. Runge, and M. A. Larson. 2011. Incorporating parametric uncertainty into population viability analysis models. Biological Conservation 144:1400–1408.

Mech, L. D. 1970. The Wolf: The Ecology and Behavior of an Endangered Species. Natural History Press, Garden City, New York.

Mech, L. D. 1995. The Challenge and Opportunity of Recovering Wolf Populations. Conservation Biology 9:270–278.

Mech, L. D., S. M. Goyal, W. J. Paul, and W. E. Newton. 2008. DEMOGRAPHIC EFFECTS OF CANINE PARVOVIRUS ON A FREE-RANGING WOLF POPULATION OVER 30 YEARS. Journal of Wildlife Diseases 44:824–836.

Mills, L. S. 2013. Conservation of wildlife populations: demography, genetics and management. Second edition.

Mitchell, M. S., J. A. Gude, K. Podruzny, E. E. Bangs, J. Hayden, R. M. Inman, M. D. Jimenez, Q. Kujala, D. H. Pletscher, and J. Rachael. 2016. Management of wolves in the US Northern Rocky Mountains is based on sound science and policy. Science. <http://science.sciencemag.org/content/350/6267/1473/tab-e-letters>.

Montana Fish, Wildlife and Parks. 2022. Wolf and Furbearer Trapping and Hunting Regulations. <https://fwp.mt.gov/binaries/content/assets/fwp/hunt/regulations/2022/wolf-and-furbearer-final-for-web.pdf.> Accessed 23 March 2023.

Morris, W. F., and D. F. Doak. 2002. Quantitative Conservation Biology: Theory and Practice of Population Viability Analysis. Sinauer Associates, Inc. Publishers, Sunderland, MA.

Murray, D. L., D. W. Smith, E. E. Bangs, C. Mack, J. K. Oakleaf, J. Fontaine, D. Boyd, M. Jiminez, C. Niemeyer, T. J. Meier, D. Stahler, J. Holyan, and V. J. Asher. 2010. Death from anthropogenic causes is partially compensatory in recovering wolf populations. Biological Conservation 143:2514–2524.

Nelson, B., M. Hebblewhite, V. Ezenwa, T. Shury, E. H. Merrill, P. C. Paquet, F. Schmiegelow, D. Seip, G. Skinner, and N. Webb. 2012. Prevalence of antibodies to canine parvovirus and distemper virus in wolves in the Canadian Rocky Mountains. Journal of Wildlife Diseases 48:68–76.

Petracca, L.S., B. Gardner, B.T. Maletzke, and S.J. Converse. 2024. Merging integrated population models and individual-based models to project population dynamics of recolonizing species. Biological Conservation 289: 110340.

Rhodes, J. R., C. F. Ng, D. L. de Villiers, H. J. Preece, C. A. McAlpine, and H. P. Possingham. 2011. Using integrated population modelling to quantify the implications of multiple threatening processes for a rapidly declining population. Biological Conservation 144:1081–1088.

Rodríguez-Recio, M., C. Wikenros, B. Zimmermann, and H. Sand. 2022. Rewilding by Wolf Recolonisation, Consequences for Ungulate Populations and Game Hunting. Biology 11:317.

Runge, M. C., and S. J. Converse. 2020. Introduction to risk analysis. Pages 149–155 in. Structured Decision Making: Case Studies in Natural Resource Management. Johns Hopkins University Press, Baltimore, MD.

Saunders, S. P., F. J. Cuthbert, and E. F. Zipkin. 2018. Evaluating population viability and efficacy of conservation management using integrated population models. Journal of Applied Ecology 55:1380–1392.

Seddon, P. J., and D. P. Armstrong. 2016. Reintroduction and Other Conservation Translocations: History And Future Developments. Pages 7–28 in. Reintroduction of Fish and Wildlife Populations. University of California Press.

Seddon, P. J., D. P. Armstrong, and R. F. Maloney. 2007. Developing the Science of Reintroduction Biology. Conservation Biology 21:303–312.

Shivik, J. A., A. Treves, and P. Callahan. 2003. Nonlethal Techniques for Managing Predation: Primary and Secondary Repellents. Conservation Biology 17:1531–1537.

Smith, D. W., E. E. Bangs, J. K. Oakleaf, C. Mack, J. Fontaine, D. Boyd, M. Jimenez, D. H. Pletscher, C. C. Niemeyer, T. J. Meier, D. R. Stahler, J. Holyan, V. J. Asher, and D. L. Murray. 2010. Survival of Colonizing Wolves in the Northern Rocky Mountains of the United States, 1982-2004. Journal of Wildlife Management 74:620–634.

Stronen, A. V., G. J. Forbes, P. C. Paquet, G. Goulet, T. Sallows, and M. Musiani. 2012. Dispersal in a plain landscape: short-distance genetic differentiation in southwestern Manitoba wolves, Canada. Conservation Genetics 13:359–371.

Treves, A., and K. U. Karanth. 2003. Human-Carnivore Conflict and Perspectives on Carnivore Management Worldwide. Conservation Biology 17:1491–1499.

Vilà, C., A. Sundqvist, Ø. Flagstad, J. Seddon, S. B. rnerfeldt, I. Kojola, A. Casulli, H. Sand, P. Wabakken, and H. Ellegren. 2003. Rescue of a severely bottlenecked wolf (Canis lupus) population by a single immigrant. Proceedings of the Royal Society of London. Series B: Biological Sciences 270:91–97.

Wabakken, P., H. Sand, O. Liberg, and A. Bjärvall. 2011. The recovery, distribution, and population dynamics of wolves on the Scandinavian peninsula, 1978-1998. Canadian Journal of Zoology 79: 710–725.

WDFW. 2020. Wolf-livestock interaction protocol. <https://wdfw.wa.gov/sites/default/files/2020-09/20200915_wdfw_wolf_livestock_interaction_protocol.pdf>. Accessed 23 March 2023.

WDFW et al. 2021. Washington Gray Wolf Conservation and Management 2020 Annual Report. <https://wdfw.wa.gov/sites/default/files/publications/02256/wdfw02256.pdf.> Accessed 23 March 2023.

Wiles, G. J., H. L. Allen, and G. E. Hayes. 2011. Wolf conservation and management plan. <https://wdfw.wa.gov/sites/default/files/publications/00001/wdfw00001.pdf.> Accessed 23 March 2023.

