## Appendix for "Forecasting dynamics of a recolonizing wolf population under different management strategies"

**Appendix S1. Results of all scenarios related to management actions and system uncertainty**

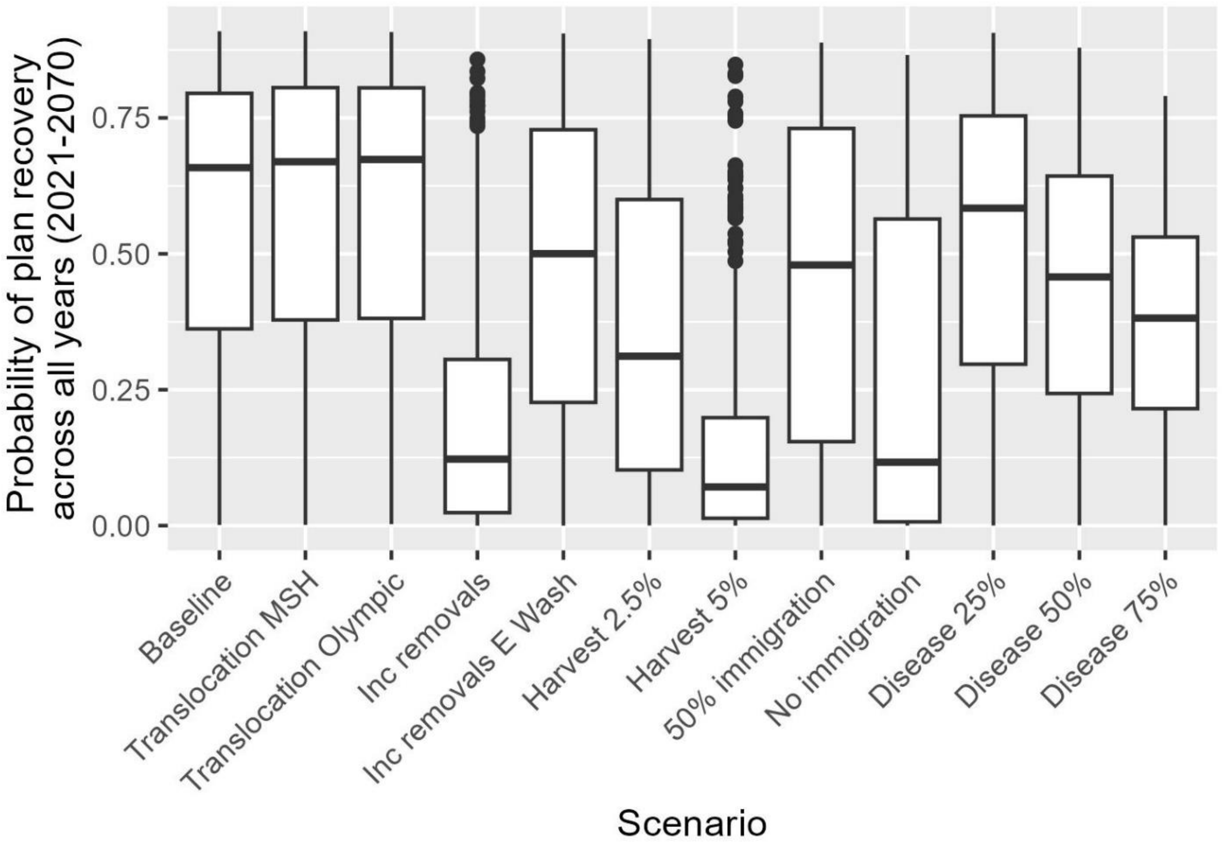


Figure S1. Probability of meeting plan recovery across all years (2021-2070) for 12 scenarios related to management and system uncertainty. Plan recovery is considered having four breeding pairs in each recovery region, with three additional breeding pairs anywhere in the state. The center line represents the median and boxes represent the 50% prediction interval. The points represent individual data points (from 50,000 samples) that are outliers.

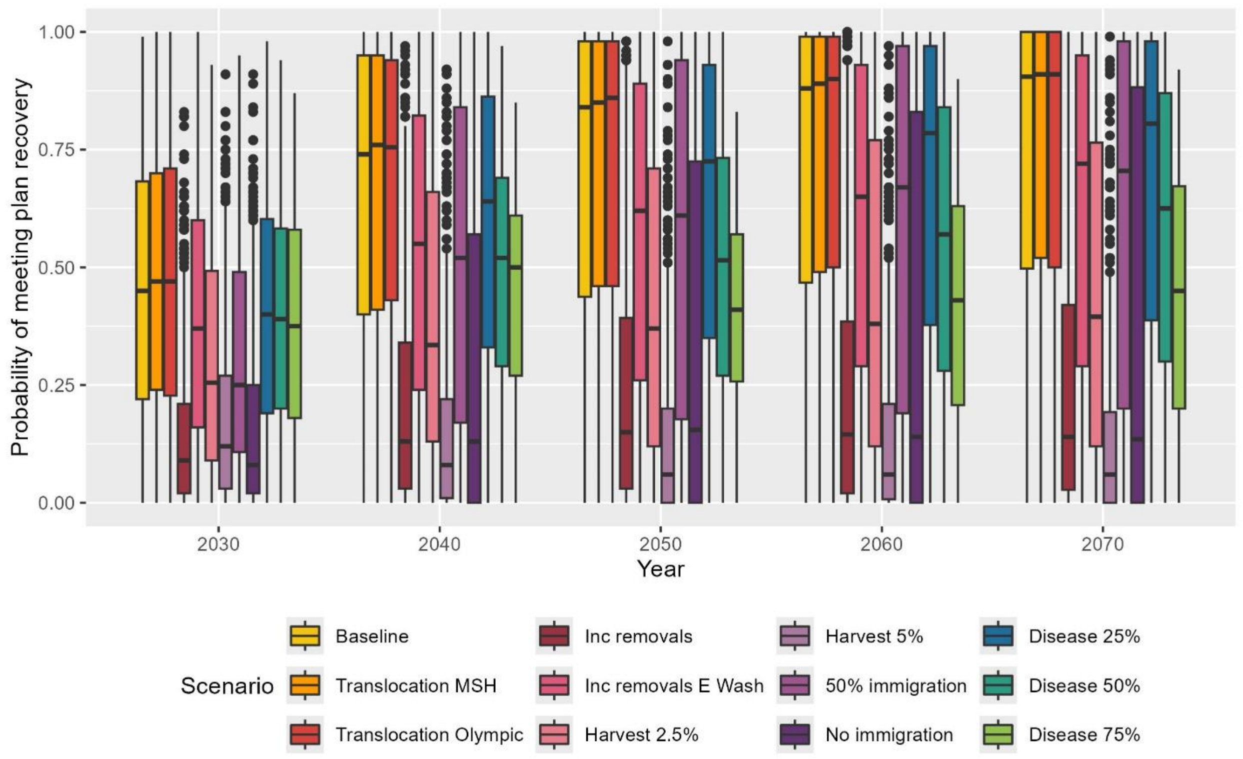


Figure S2. Probability of meeting plan recovery at various time points over the period 2021-2070 for 12 scenarios related to management and system uncertainty. Plan recovery is considered having four breeding pairs in each recovery region, with three additional breeding pairs anywhere in the state. The center line represents the median and boxes represent the 50% prediction interval. The points represent individual data points (from 50,000 samples) that are outliers.

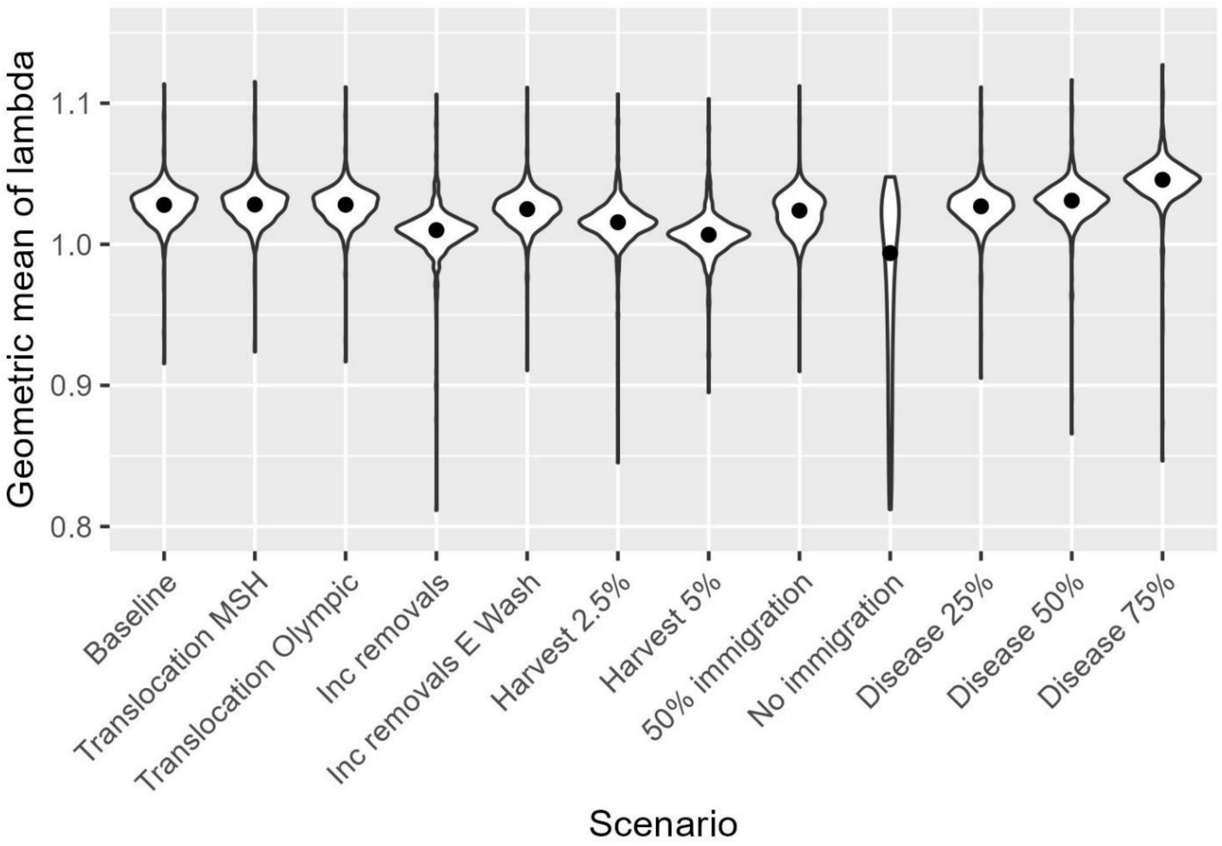


Figure S3. Geometric mean of lambda of the gray wolf (*Canis lupus*) population in Washington State, USA, over the time period 2021-2070, for 12 scenarios related to management and system uncertainty.

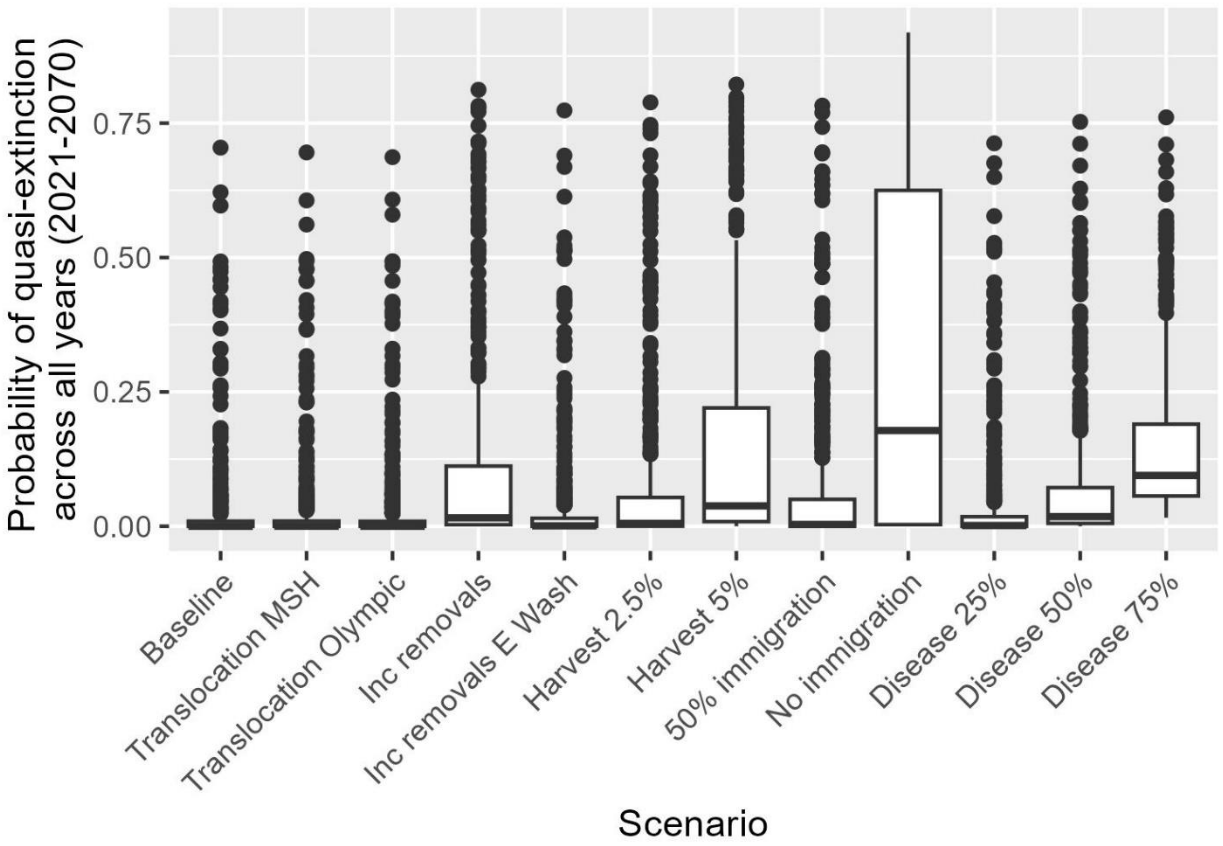


Figure S4. Probability of quasi-extinction across all years (2021-2070) for 12 scenarios related to management and system uncertainty. Quasi-extinction occurs when there are <92 adult wolves in the state and <24 adult wolves in each recovery region. The center line represents the median and boxes represent the 50% prediction interval. The points represent individual data points (from 50,000 samples) that are outliers.

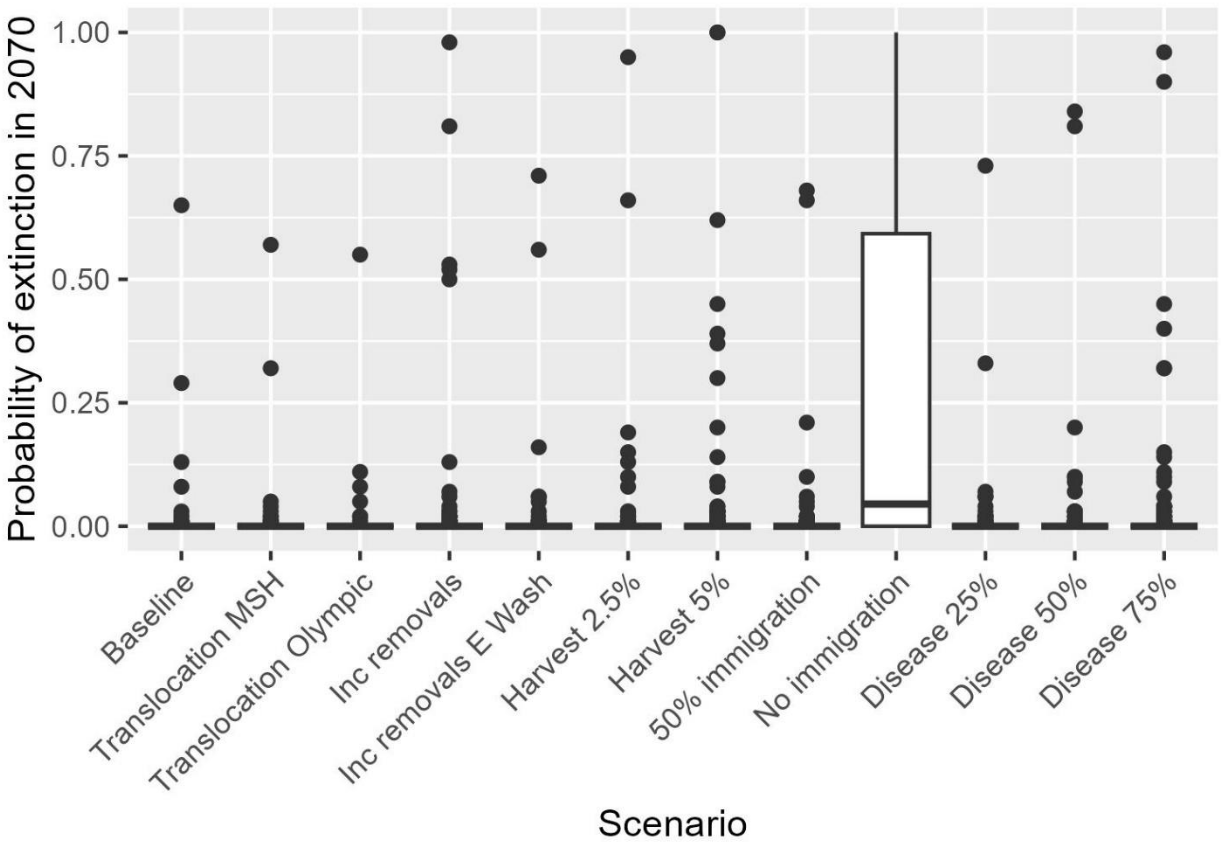


Figure S5. Probability of extinction in the year 2070 for 12 scenarios related to management and system uncertainty. Extinction occurs when there are zero wolves left in Washington State. The center line represents the median and boxes represent the 50% prediction interval. The points represent individual data points (from 50,000 samples) that are outliers.

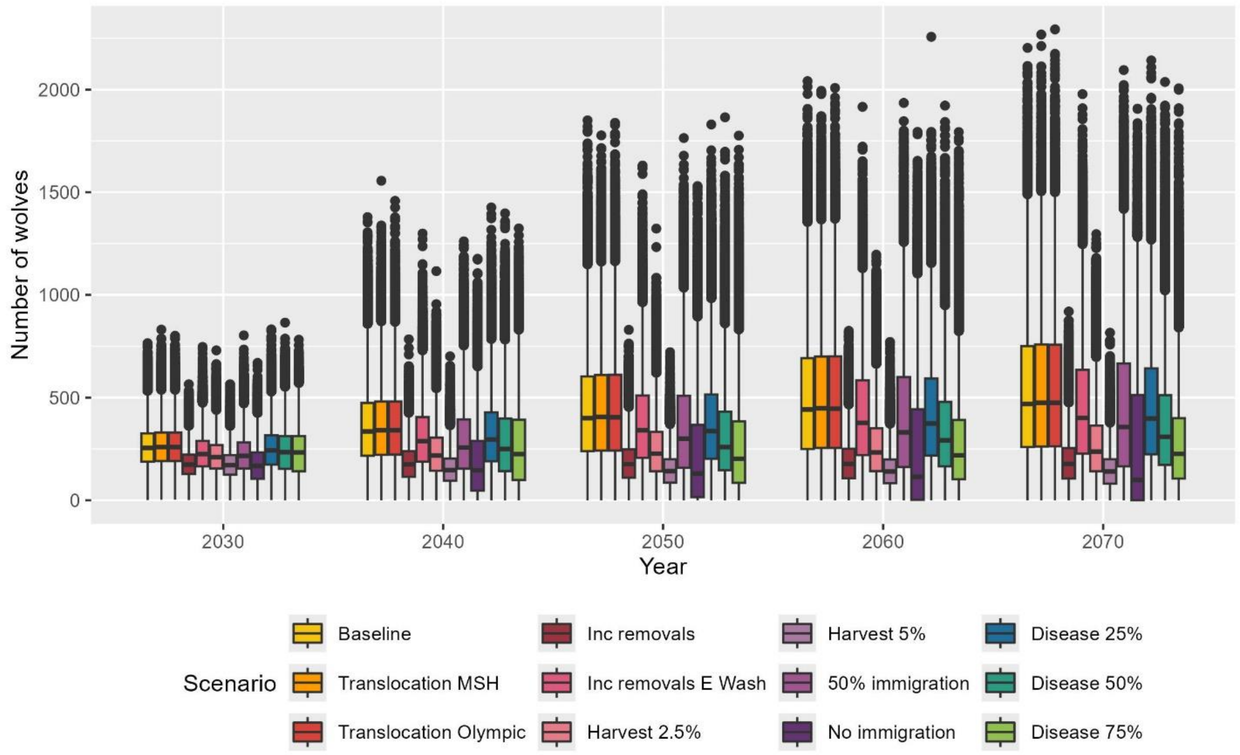


Figure S6. Estimated number of wolves in Washington State, USA, for 12 scenarios related to management and system uncertainty. Black line indicates the median and boxes represent the 50% prediction interval. The points represent individual data points (from 50,000 samples) that are outliers.
